# Live imaging of chromatin distribution in muscle nuclei reveals novel principles of nuclear architecture and chromatin compartmentalization

**DOI:** 10.1101/2020.06.21.163360

**Authors:** Daria Amiad-Pavlov, Dana Lorber, Gaurav Bajpai, Samuel Safran, Talila Volk

## Abstract

Packaging of the chromatin within the nucleus serves as an important factor in the regulation of transcriptional output. However, information on chromatin architecture on nuclear scale in fully differentiated cells, under physiological conditions and in live organisms, is largely unavailable. Here, we imaged nuclei and chromatin in muscle fibers of live, intact *Drosophila* larvae. In contrast to the common view that chromatin is distributed throughout the nuclear volume, we show that the entire chromatin, including active and repressed regions, forms a peripheral layer underneath the nuclear lamina, leaving a chromatin-devoid compartment at the nucleus center. Importantly, visualization of nuclear compartmentalization required imaging of un-fixed nuclei embedded within their intrinsic tissue environment, with preserved nuclear volume. Upon fixation of similar muscle nuclei, we observed an average of three-fold reduction in nuclear volume caused by dehydration and evidenced by nuclear flattening. In these conditions, the peripheral chromatin layer was not detected anymore, demonstrating the importance of preserving native biophysical tissue environment. We further show that nuclear compartmentalization is sensitive to the levels of lamin C, since over-expression of lamin C-GFP in muscle nuclei resulted in detachment of the peripheral chromatin layer from the lamina and its collapse into the nuclear center. Computer simulations of chromatin distribution recapitulated the peripheral chromatin organization observed experimentally, when binding of lamina associated domains (LADs) was incorporated with chromatin self-attractive interactions. Reducing the number of LADs led to collapse of the chromatin, similarly to our observations following lamin C over-expression. Taken together, our findings reveal a novel mode of mesoscale organization of chromatin within the nucleus in a live organism, in which the chromatin forms a peripheral layer separated from the nuclear interior. This architecture may be essential for robust transcriptional regulation in fully differentiated cells.

## Introduction

While three-dimensional (3D) organization of the genome has been directly linked to gene regulation, information regarding nuclear-scale 3D chromatin organization under native physiological conditions is limited. Recent advances in imaging, sequencing and modeling approaches have greatly enhanced our understanding of 3D genome organization, bridging the gap between single-cell spatial information and genome-wide linear sequence interactions ^1^. However, the absence of experiments that visualize the global 3D chromatin organization within the native tissue environment limits the interpretation of current data.

The genome is packed within the nucleus in a non-random manner and organized in a hierarchical manner, with several characteristic length scales. The basic nucleosome unit consists of DNA wrapped around a core octamer of histone proteins H2A, H2B, H3, and H4. Nucleosomes are organized into 10-nm “beads on a string” fiber that together with additional proteins define the chromatin. While the textbook view suggests hierarchical higher-order folding of chromatin fibers (10 nm, 30 nm, 100 nm, and higher), recent advanced imaging under more physiological conditions challenged this view, describing a more complex and heterogeneous chromatin organization into clutches and domains of nucleosomes of varying sizes and densities ^1–4^. On the nuclear scale, a range of imaging approaches, as well as Hi-C analysis, suggest that chromatin of an interphase nucleus is partitioned into distinct chromosomal territories several µm long, as was documented in various mammalian and *Drosophila* cells ^5–8^. A functional model for global chromatin organization suggests that chromatin territories occupied by the chromosomes are pervaded by a channel network devoid of chromatin, termed inter-chromatin compartments ^9,10^.

Several lines of evidence suggest a non-uniform distribution of chromatin as a function of the radial distance from the center of the nucleus. Interphase chromosomes and specific gene loci within chromosomes have preferential radial positions, with gene rich chromosomes positioned towards the nucleus interior. However, such radial positioning is tissue specific and highly variable among cells ^7,11^. Furthermore, the nuclear lamina, a thick meshwork of intermediate filaments associated with the inner nuclear membrane, is a major regulator of chromatin architecture, as it tethers mostly dense heterochromatin at specific sequences termed lamina-associated domains (LADs). Consistently, genome-wide DamID analysis performed with lamin B or lamin A in culture conditions identified LADs mostly as gene poor and transcriptionally silenced sequences ^12^. Specifically, variations in lamin A/C levels, induced by either its up-or down regulation, or as observed in laminopathy-associated mutations in lamin A/C, drive heterochromatin detachment from the nuclear lamina ^13,14^. This was accompanied by chromatin de-condensation and reduced levels of HP1 and H3K9me3 repressive marks in mammals ^15,16^, *C. elegans* ^17^ as well as in *Drosophila* cells ^18,19^. Taken together, these findings suggested that LADs are specifically sensitive to the levels of lamin A/C at the nuclear lamina.

Chromatin is estimated to fill about 15-60% of the nuclear volume ^7,9^, and shares the nucleoplasm with other dense, membrane-less organelles such as the nucleolus and nuclear speckles. Furthermore, imaging of either fixed nuclei, live cells in culture, or cells isolated from their native tissue environment suggests that chromatin is distributed throughout the entire nuclear space. However, the lack of information on nuclei within their intrinsic tissue environment, where cells and nuclei adopt a specific 3D morphology, obscures the conclusions regarding the actual chromatin distribution in mature, differentiated cells. Chromatin organization can change in space and time in a physiological, adaptive manner, where nuclear morphology is an important contributor that reflects the balance between cytoplasmic forces acting on the nucleus and the collective mechanical resistance of the lamina and the chromatin ^20–23^. Additionally, nuclear morphology and chromatin organization can be altered due to experimental methodologies such as reduction in cell and nucleus volume due to fixation, change in osmolality, or as a result of culture on surfaces of various stiffness conditions ^24–26^. Recent study that analyzed live telomere dynamics in the liver *in vivo*, reported significantly different dynamics from that observed in cultured cells ^27,28^. It is therefore critical to reveal chromatin organization in its tissue intrinsic physiological conditions.

Here we combined a custom-made device placed on top of a confocal microscope stage ^29^, with genetic labeling of the chromatin and the nuclear lamina, to characterize *in vivo* nuclear scale 3D chromatin organization in fully differentiated muscle of live, intact *Drosophila* larva. Imaging of live 3D chromatin organization was reported for various cell lines ^2,30,31^, as well as in the isolated *Drosophila* larva salivary gland ^32^. We present here the first 3D analysis of global chromatin organization of fully differentiated nuclei, with preserved physiologic environment, within a live, intact organism. Our live imaging demonstrates significantly different nuclear dimensions and higher nuclear volume compared to fixed preparations ^33,34^. Importantly, the preserved ellipsoid shape and volume of the live nuclei enabled visualization of a novel, nuclear-scale mode of global chromatin organization, where both active and non-active chromatin regions are distributed at the periphery of the muscle nucleus, forming a substantial region that is devoid of chromatin in the nucleus interior. This peripheral chromatin organization was sensitive to the levels of lamin A/C, since over-expression of lamin A/C resulted in chromatin detachment from the nuclear lamina, leading to a condensation of chromatin towards the center of the nucleus. We further present simulation results of a polymer model, demonstrating that a balance between chromatin association with the lamina, and substantial, effective chromatin self-attraction can explain the global peripheral chromatin distribution that we observed experimentally. Reduced LAD attachment to the lamina in the simulations resulted in chromatin collapse towards the center, similar to the experimental observation of lamin A/C over-expression in muscle nuclei. Our results reveal a novel mode of nuclear organization in fully differentiated cells embedded within their native tissue environment, which have peripheral chromatin-rich and central chromatin-devoid regions, and demonstrate the sensitivity of nuclear-scale chromatin organization to changes in the lamina composition, as well as altered nuclear shape and volume.

## Results

### Live imaging of 3D chromatin distribution in intact *Drosophila* larval muscles reveals novel, nuclear-scale chromatin organization

The current view of higher-order chromatin organization depicts chromatin as distributed throughout the entire nuclear volume ^7^. So far, most of the studies describing 3D chromatin organization were conducted either on cultured cells, or on cells in fixed tissue preparations ^3,8,19,21,35^. To study 3D chromatin organization under physiological conditions in a live organism, we have developed a setup to visualize nuclei and chromatin in live, intact *Drosophila* larva. This setup is based on a minimally constrained device designed for imaging stationary or contractile muscles and their nuclei in an intact *Drosophila* larva, preserving tissue and nuclear native environment ^29^. Figure 1A shows a 3D view of a single muscle nucleus, cut through the middle, of live, intact 3^rd^ instar *Drosophila* larva. The nuclear envelope was labeled with UAS-nesprin/klar-GFP driven by a muscle-specific driver (*Mef2-Gal4*), and chromatin was labeled by ubiquitous expression of His2B-mRFP, a common live mark for total chromatin ^2,8,30,31^.

**Figure 1:**
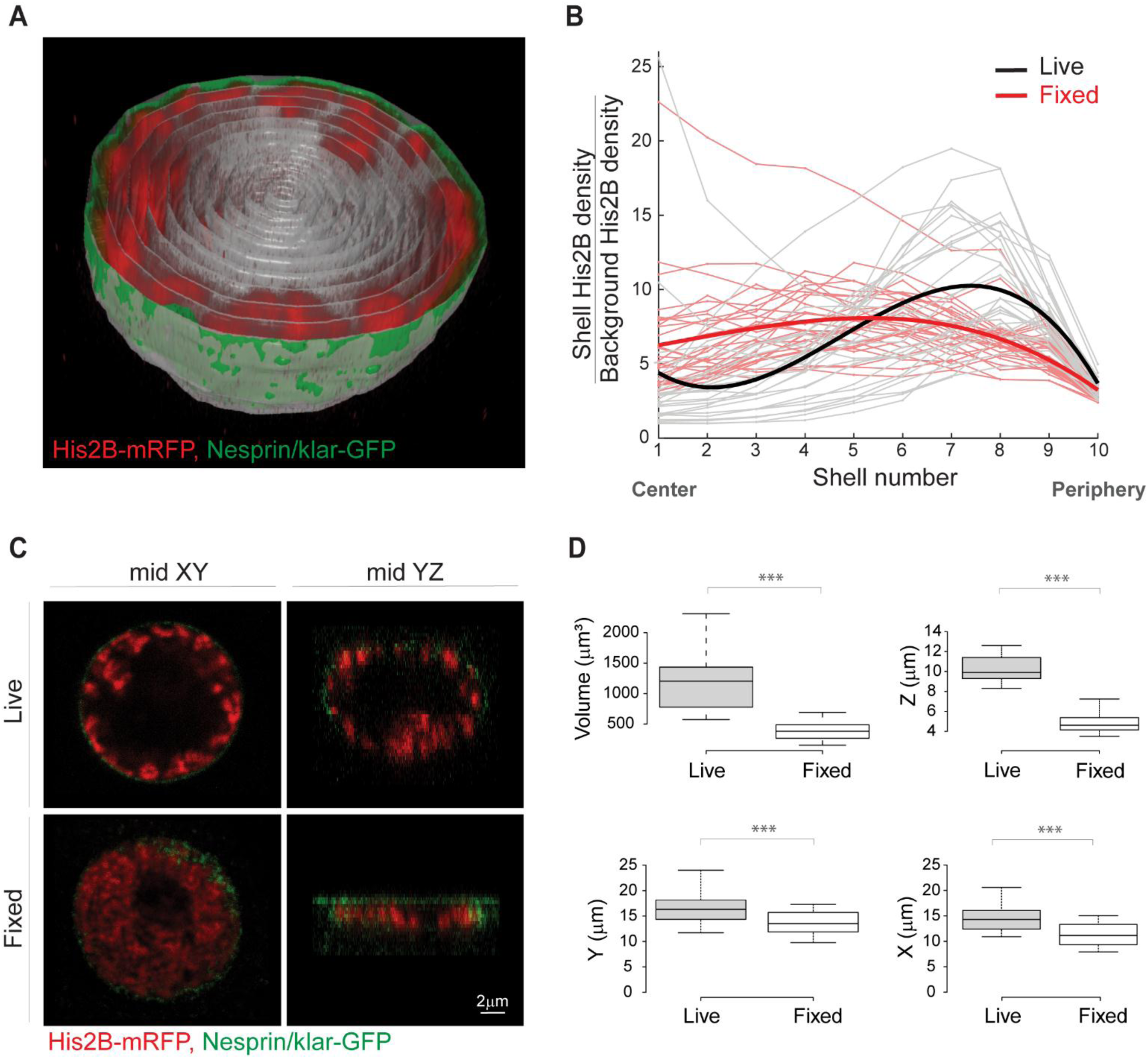
Peripheral chromatin organization in live *Drosophila* larva muscle nuclei, not detected in fixed nuclei. (A) 3D view of a single live muscle nucleus cut through the middle. Chromatin is labeled with His2B-mRFP (red) and nuclear envelope with Nesprin/klar-GFP (green). For quantification of radial chromatin distribution, the segmented nucleus is divided into 10 concentric 3D radial shells (gray). (B) His2B fluorescent signal density for each shell, normalized by the background signal density, is plotted from center to periphery. The resulting radial chromatin distribution profile demonstrates peripheral trend in live nuclei (individual nuclei in gray, fitted average in bold black, N=27), which is significantly different from the more uniform radial chromatin distribution profile in fixed preparations (individual nuclei in light red, fitted average in bold red, N=30; p<0.001). (C) Mid confocal XY plane of a representative nucleus from live, intact larva shows that chromatin is organized at the periphery of the nucleus (top left), whereas in a representative nucleus from a fixed preparation, chromatin is expanded into most of the nuclear space (bottom left). Mid YZ plane of the live nucleus demonstrates preserved ellipsoid shape and peripheral chromatin (top right), whereas the mid YZ plane from the fixed nucleus reveals substantial flattening and altered chromatin organization (bottom right). (D) Significant reduction (p<0.001) in nuclear volume, as well as in the short, middle, and long nuclear dimensions in fixed preparations (N=30) compared to live (N=27).

Remarkably, 3D visualization of the live muscle nuclei revealed a peripheral distribution of the chromatin, with a substantial region in the interior of the nucleus which was devoid of chromatin (Movie S1). To quantify chromatin distribution along the radial direction, each nucleus was segmented in 3D and divided into 10 concentric shells (Figure 1A, gray traces). Chromatin density for each shell was calculated from the sum of His2B-mRFP fluorescent intensity, divided by the shell volume. The background His2B-mRFP density was calculated from an additional shell outside of the nuclear envelope. For each nucleus we generated a radial chromatin distribution profile, by plotting the chromatin fluorescent density, normalized by the background fluorescent density, from the center to the periphery of the nucleus (Figure 1B); Note that shell 10 was defined as the outer nuclear envelope. To study the robustness of the phenomena, we analyzed the radial chromatin distribution profiles of nuclei from at least 3 different, randomly chosen muscles along the larva length, and from 5 different larvae (gray traces in Figure 1B, N=27). Mean radial chromatin distribution demonstrates a robust peripheral profile for the live nuclei (bold black in Figure 1B, fitted with 3^rd^ order polynomial, linear mixed effect model). Note that a few individual live nuclei also have high chromatin density in the center, in addition to the high peripheral chromatin density. Examination of these few nuclei revealed a chromatin branch extended from one side of the nucleus to the other, but still most of the nucleus center is devoid of chromatin, as demonstrated in Figure S1.

In contrast to the peripheral chromatin distribution we observed in muscles of live larvae, previous studies from our group and of others, performed on muscle nuclei from fixed *Drosophila* larva, depicted chromatin that is fairly uniformly distributed throughout the entire nuclear volume ^33,34^. To investigate this discrepancy, we compared chromatin distribution in muscle nuclei visualized in the live larvae, with that observed in fixed larval preparations, utilizing identical genotype and method of chromatin labeling and similar imaging setup and image analysis. Notably, the radial chromatin distribution profile of fixed nuclei (Figure 1B, individual nuclei in light red, N=30, fitted average in bold red) was significantly different from that of the live larvae (black, p<0.001), indicating a relatively homogenous chromatin distribution throughout the nuclear volume (Figure 1C).

To reveal possible explanations for the marked difference in global chromatin organization between live and fixed nuclei, we measured nuclear dimensions and volume of live and fixed nuclei. Figure 1C shows a single mid-section in the XY plane (left) and a single mid-section in the XZ plane (right) of a confocal Z-stack taken from live (top) or fixed (bottom) nuclei of comparable size. In the XY plane, chromatin distribution of the live nucleus appeared peripheral, whereas in the fixed nucleus, the chromatin distributed throughout most of the nuclear volume, excluding only the region occupied by the nucleolus. Importantly, whereas the live nucleus preserved its volume and ellipsoidal shape, the fixed nucleus exhibited substantial flattening, forming a disk-like shape ^34^, as depicted in the XZ plain. Figure 1D summarizes the reduction in nuclear volume and nuclear dimensions upon fixation. On average, there was a 3.1-fold decrease in nuclear volume from 1183.1 µm^3^ in live nuclei to 380.8 µm^3^ in fixed nuclei (p<0.001), mostly due to a 2.1-fold reduction in the Z diameter, from an average of 10.3 µm in live nuclei to 4.9 µm in fixed nuclei (p<0.001). Note that the Z dimension is perpendicular to the muscle fiber axis. There was also a smaller, but significant 1.3-fold decrease in the X and Y diameters of fixed nuclei, from 14.4 µm and 16.3 µm in live nuclei to 11.2 µm and 13.7 µm in the fixed nuclei, respectively (p<0.001).

It has been reported that tissue fixation can cause cellular and nuclear volume reduction due to dehydration of the sample ^25,36^. We therefore speculate that *Drosophila* larva muscle nuclei appeared flat in fixed preparations due to dehydration. We hypothesize that our live, intact organism setup preserved the physiological conditions and the native environment of the muscle tissue, resulting in nuclei with more spherical rather than flattened shape. Importantly, the spherical shape of the nuclei observed in the live setup allowed us to distinguish the chromatin-containing compartment localized at the periphery of the nucleus from the chromatin-devoid compartment in the interior of the nucleus. This higher-order chromatin organization is not observed in fixed conditions, in which muscle nuclei are flattened due to the decrease in the nuclear Z-dimension, and might be undetectable in other studies of interphase nuclei where the native nuclear volume and dimensions are often compromised due to fixation or cell culture conditions. Interestingly, similar peripheral chromatin organization was observed in nuclei of fixed 8-cell stage bovine embryo obtained by *in vitro* fertilization (IVF) and was reported to shift into a conventional distribution (chromatin expanded throughout the nuclear volume) with development progression into 20-cell embryo ^10^. Importantly, the study reported substantial reduction in nuclear volume upon developmental stage advancement, further supporting the link between nuclear volume and global chromatin organization as suggested by our results.

To further support the observation of peripheral chromatin organization in the live, intact setup, we utilized His2Av-GFP as an additional, commonly used marker for total chromatin labeling ^32^. Figure 2A shows a mid XY plane of a nucleus from live larva muscle, co-labeled with His2Av-GFP (A), and His2B-mRFP(A’). The merged view (A’’) indicates high colocalization of both histone tags, providing additional support to the hypothesis that the observed peripheral chromatin distribution is independent of the labeling type.

**Figure 2:**
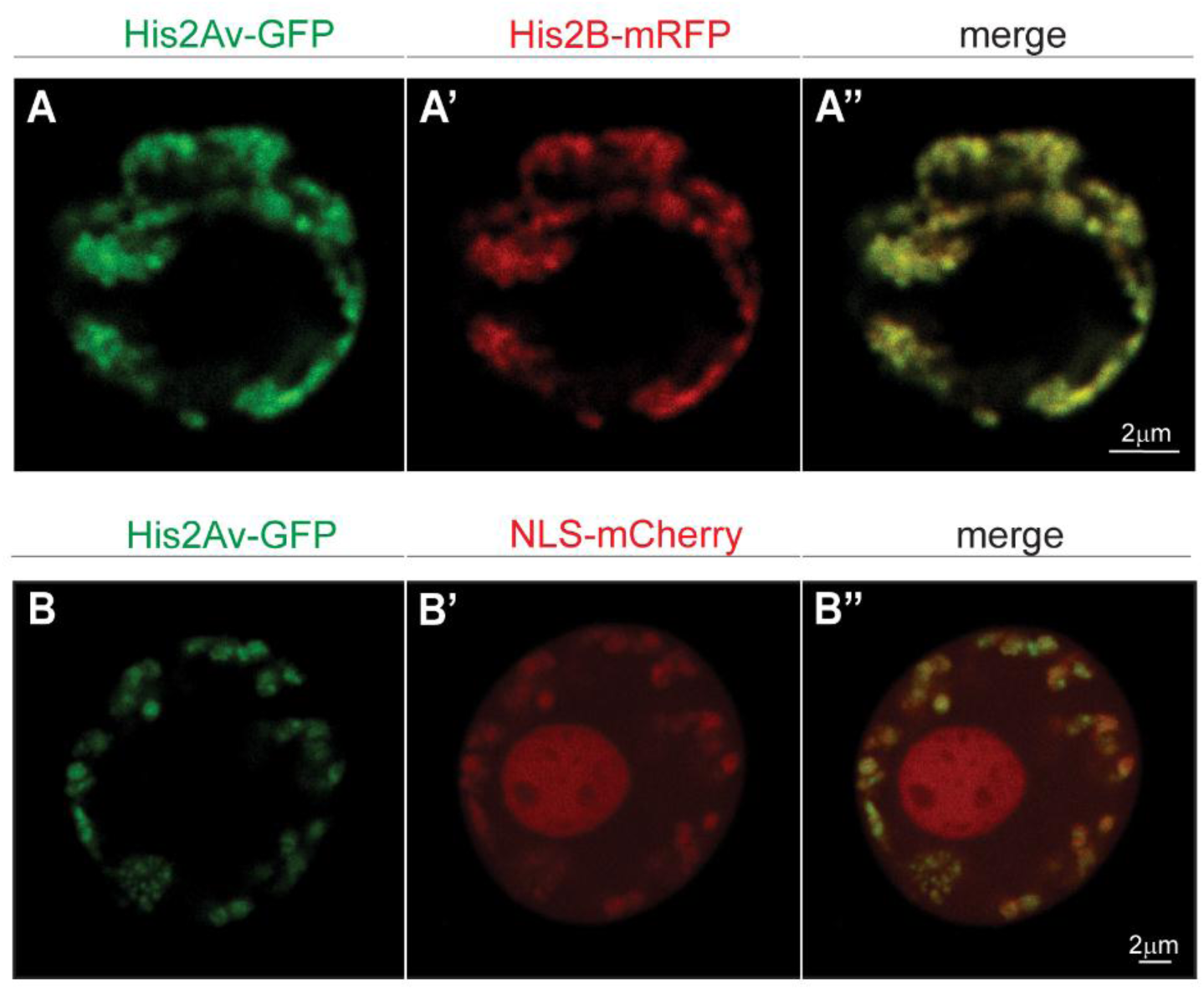
Peripheral chromatin organization in the live nucleus is also observed by His2Av labeling, and the chromatin devoid compartment does not correspond to the nucleolus. (A-A’’) Co-labeling of a live muscle nucleus with two independent histone labels, His2Av-GFP (A), and His2B-mRFP (A’), shows high co-localization (A’’), supporting peripheral chromatin organization. (B-B’’) live muscle nucleus co-labeled with His2Av-GFP (B) and NLS-mCherry (B’) demonstrates the nucleolus, as well as an additional chromatin devoid volume in the nucleus interior.

The nucleolus is a membrane-less, phase separated organelle within the nucleus. Each nucleus of the *Drosophila* larval muscle has a single nucleolus estimated to occupy roughly 20% of the projected nucleus area ^34^. To address whether a large nucleolus compartment might exclude chromatin from the central region of the nucleus, we expressed mCherry fused to nuclear localization signal (NLS) in muscles, together with His2Av-GFP (Figure 2B and 2B’, respectively). The NLS-mCherry appeared throughout the entire nucleoplasm with a denser accumulation inside the nucleolus, which was verified by co-labeling with fibrillarin (Figure S2). The merged view in Figure 2B’’ demonstrates a large chromatin-devoid region in the nucleus interior, that did not overlap the nucleolus domain. Together, these results suggest that in live *Drosophila* larval muscles, chromatin is arranged at the periphery of the nucleus, with a substantial region devoid of chromatin in the nucleus interior. Interestingly, we found that the peripheral organization of chromatin was not unique to the muscle tissue and was also observed in larval epidermal nuclei (Movie S2).

### Chromatin localized at the nuclear periphery contains active chromatin regions

A non-random higher order chromatin organization is a hallmark of the interphase nucleus 3D structure ^9,13^. With respect to the radial distribution, it is well established that the nuclear periphery is enriched with condensed heterochromatic regions of the genome, and that LADs, representing DNA sequences that interact with the nuclear lamina, are largely transcriptionally repressive ^11,12,37^. The nucleus interior on the other hand, has been reported to contain gene-rich chromatin, which is often actively transcribed ^7^. To rule out the possibility that the peripherally organized chromatin observed in our study represents mostly dense heterochromatin, while more open, active chromatin in the nucleus interior was undetected by our imaging approach, we co-labeled nuclei with a live tag for active chromatin. Sato et al. have developed a fluorescently-labeled modification-specific antibody (mintbody), by fusing EGFP to a single-chain variable fragment antibody with high specificity to H3K9ac, which allows *in vivo* tracking of active chromatin domains with minimal interference to cell function ^38^. Muscle nuclei of live *Drosophila* larvae co-labeled with H3K9ac-EGFP and His2B-mRFP were imaged as described above, to identify the active chromatin regions with respect to the peripherally distributed chromatin compartment (mid confocal planes are shown in Figure 3A and 3A’, respectively). In contrast to the His2B-mRFP signal, which shows stronger fluorescence within dense chromatin regions, the H3K9ac-EGFP is expected to exhibit higher signal intensity within the less dense, open chromatin regions. The merged image in Figure 3A’’ (and Movie S3) shows that active chromatin regions were distributed in the periphery of the nucleus, with a spatial pattern similar to that of total chromatin. The dashed boxed region of Figure 3A’’ is enlarged in Figure 3B-B’’ and highlights the inverse pattern of both labels, with respect to signal intensity. The filled white arrow points to the large, repressed chromocenter of the nucleus with a strong His2B-mRFP signal, and lacks the H3K9ac-EGFP signal of the active chromatin. The empty white arrow points to one of the active regions with diffuse chromatin indicated by weak His2B-mRFP fluorescence, but strong H3K9ac-EGFP signal intensity.

**Figure 3:**
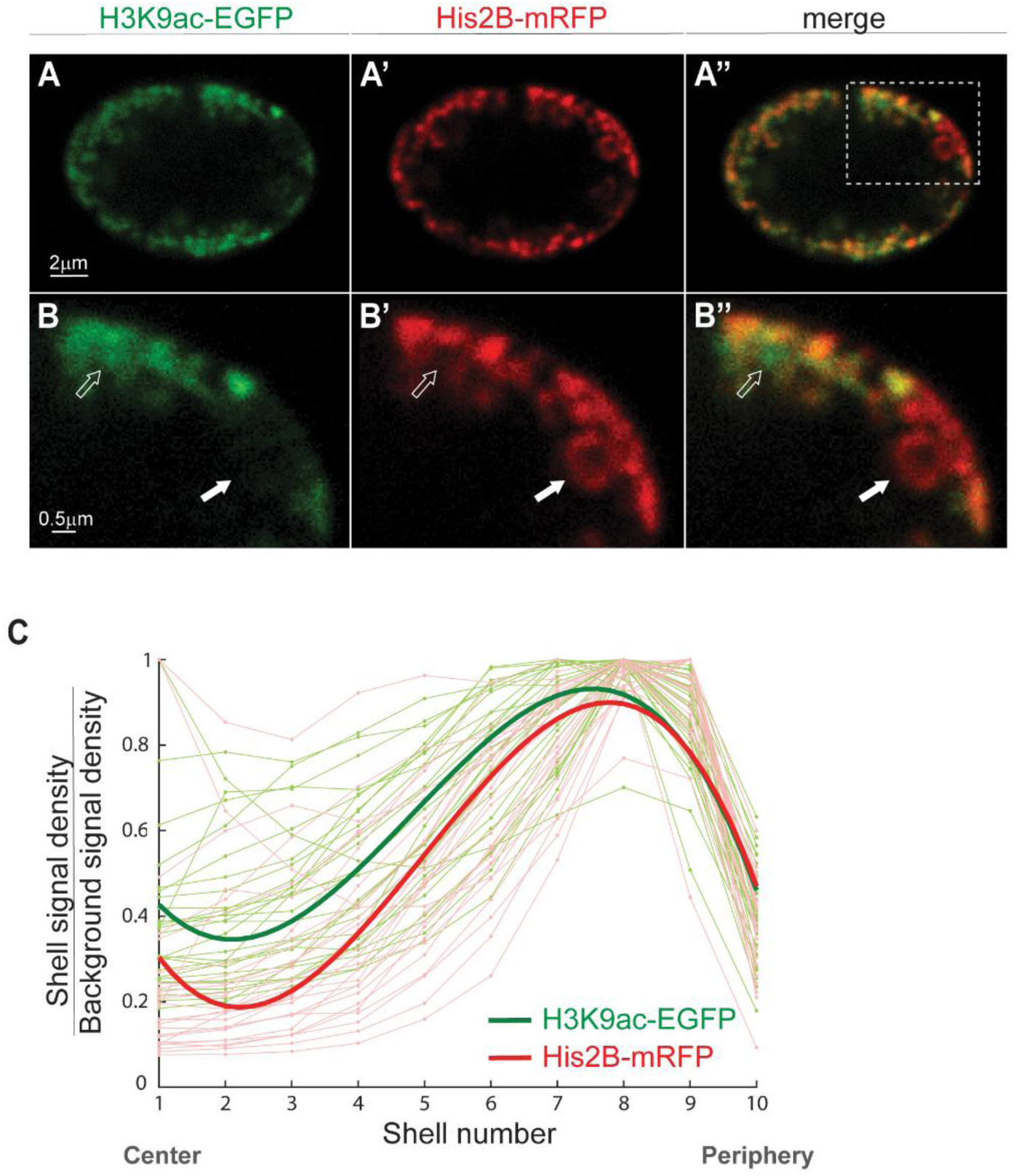
Live labeling of active chromatin regions shows spatial overlap with His2B-mRFP labeled chromatin compartment at the nuclear periphery. (A) H3K9ac-EGFP mintbody expressed in the live larva muscle nucleus demonstrates peripheral distribution of the euchromatin, similar to the His2B-mRFP spatial distribution of the total chromatin. (A’). Merged image of active and total chromatin (A’’); boxed area is enlarged in the lower panel (B-B’’). Solid white arrow points to a dense heterochromatin with strong His2B-mRFP signal (B’) and weak H3K9ac-EGFP (B). Hollow arrow points to an active region with strong H3K9ac-EGFP signal, and diffused chromatin emphasized by weak His2B-mRFP. These regions coexist in the same radial layer, but at different positions perpendicular to the radial direction. (C) H3K9ac (light green) and His2B (light red) signal density for each shell, normalized by background signal density, is plotted from center to periphery. Each light line represents a single nucleus, with signal normalized to maximum value (N=27). Bold lines represent the fitted average, with peak signal at the periphery of the nucleus for both active and overall chromatin.

Figure 3C compares the 3D radial fluorescent signal distribution profiles for the active, H3K9ac (light green) and total chromatin, His2B (light red) in individual nuclei (N=27). Each radial profile was scaled to its maximum value to simplify the comparison of the profile shapes. The fitted average radial distribution profiles indicated peripheral distribution of the active chromatin regions (bold green), similar to that of total chromatin (bold red). Statistical analysis comparing the shape descriptors of the profiles showed no significant differences between the radial distributions of the active relative to the total chromatin. The lack of a strong active chromatin signal in the nucleus interior rules out the possibility of undetectable, de-condensed chromatin, which could possibly be the case when imaged solely by His2B-mRFP labelling. Moreover, our results suggest that the active euchromatin is not separated from the heterochromatin along the radial direction, but rather separated in the perpendicular direction, into active hubs of varying sizes within the peripheral chromatin layer. This is in agreement with the observations in nuclei of fixed 8-cell bovine embryo demonstrating no radial segregation between active and repressed regions in the peripherally organized chromatin ^10^, and with a recent study showing that active chromatin forms spatially segregated clusters, mostly excluded from larger dense, heterochromatic clusters ^35^.

### Lamin C over-expression disrupts chromatin localization at the nuclear periphery

*Drosophila* is the only known invertebrate model expressing both types of the human lamins (i.e., A/C and B types), along with genome representation of additional nuclear envelope components such as the LINC complex, LEM domain, and BAF proteins. *Drosophila* lamin C, which is the homolog of mammalian A/C type lamin, displays tissue-specific expression levels in fully differentiated cells ^39,40^. Mutations in lamin A/C have been associated with aberrant chromatin organization and global detachment of chromatin from the nuclear lamina in a wide range of model organisms and tissues ^15,18,19^. Therefore, we utilized the genetic advantages of the *Drosophila* to study the effect of over-expression (OE) of lamin C on the peripheral chromatin localization that we observed in live larva muscles. Towards that end, Lamin C-GFP was expressed in muscles using the *mef2-gal4* muscle driver in combination with the ubiquitously expressed His2B-mRFP, allowing live imaging of the nuclear envelope and chromatin (Figure 4A). We did not observe in lamin C-GFP OE larvae altered development, or apparent viability phenotypes. Figure 4A (and Movie S4) shows a 3D view of a single muscle nucleus with lamin C OE, cut through the middle, in our live, intact larval setup. In agreement with previous reports, the chromatin was detached from the nuclear lamina and was condensed towards the center. The nucleus in Figure 4A is shown at a single mid XY plane in Figure 4B (nucleus 1), along with 2 additional nuclei (nucleus 2 and 3), that represent variable phenotypes of global chromatin distribution obtained following lamin C-GFP OE. This phenotypic variability was presumably caused by differential levels of the *mef2-gal4* driver in different muscles. Significantly, a general trend of disruption to the peripheral chromatin localization, and a collapse of chromatin towards the nucleus interior is demonstrated in Figure 4C showing radial His2B-mRFP density profiles for the control nuclei (gray, N=27) and nuclei with lamin C-GFP OE (light red, N=23). Each radial profile was scaled to its maximum value to aid with shape comparison. Despite the variability in radial chromatin distribution within the lamin C-GFP OE group, the fitted average radial density profile (bold red) demonstrates a substantial shift in chromatin density from the periphery towards the center, compared with control nuclei (bold black). Statistical analysis comparing the shape descriptors of control and lamin C-GFP OE profiles showed significant differences in their radial chromatin distribution (p<0.001).

**Figure 4:**
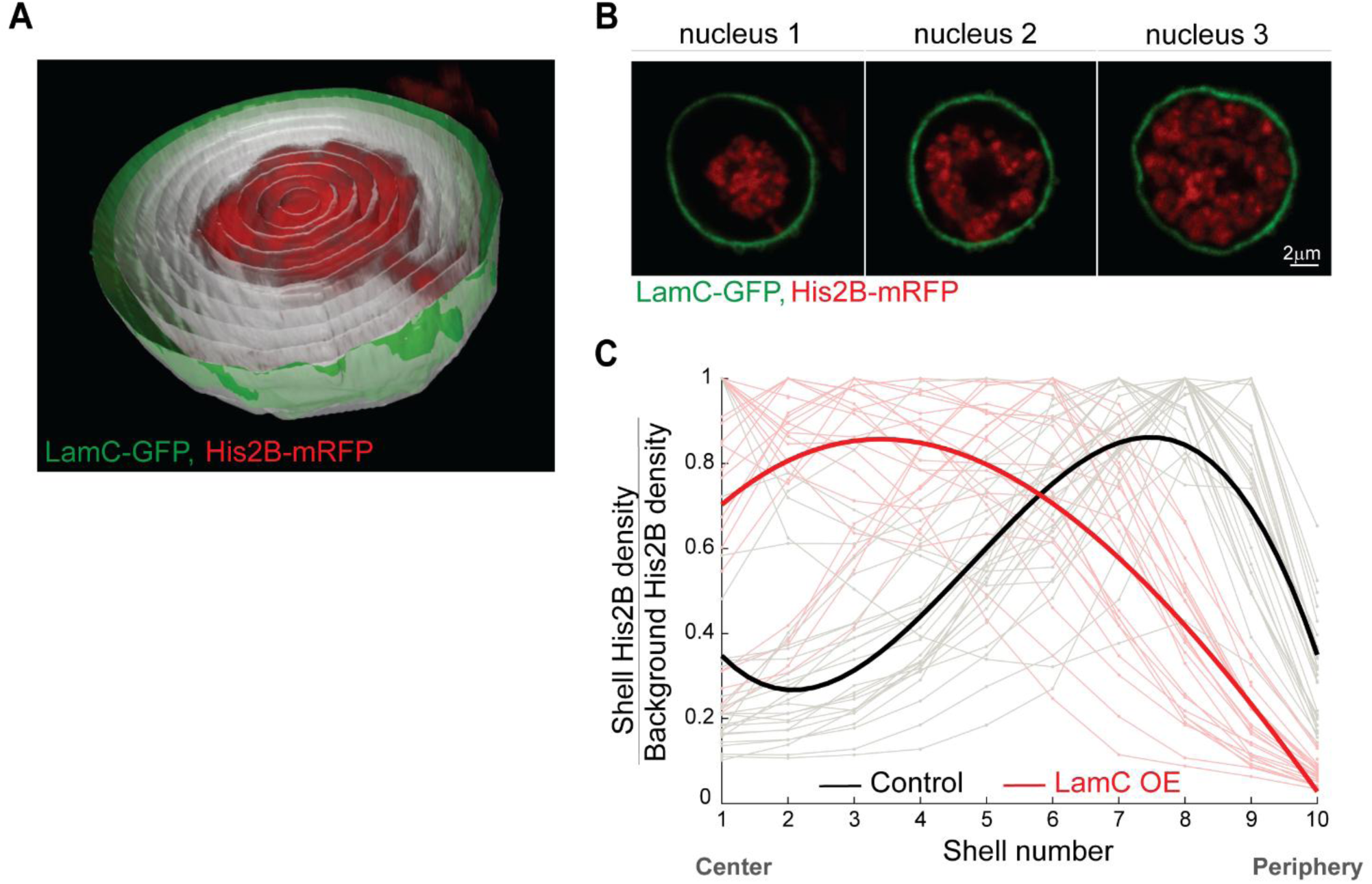
Lamin A/C over-expression in the Drosophila larva muscle disrupts peripheral chromatin localization, driving chromatin condensation towards the center of the nucleus. (A) 3D view of a live nucleus, over-expressing lamin C, cut through the middle. Chromatin is labeled with His2B-mRFP and nuclear envelope with lamin C-GFP. The segmented nucleus is divided into 10 concentric 3D radial shells (gray) for quantification of radial chromatin distribution. (B) Variable phenotypes of chromatin distribution in lamin C OE nuclei from different muscles. (C) His2B signal density for each shell, normalized by background signal density, is plotted from center to periphery. Each individual line was normalized to maximum value. The resulting radial chromatin density profile demonstrates the trend towards central localization in the live lamin C OE group (individual nuclei in light red, fitted average in bold red, N=23), compared with the peripheral trend in live control nuclei (individual nuclei in gray, fitted average in bold black, N=27).

Taken together, our results demonstrate that elevated lamin C levels in the fully differentiated larva muscle caused detachment of chromatin from the lamina and loss of peripheral organization, in agreement with previous studies demonstrating that both up and down regulation of lamin C levels lead to disrupted 3D chromatin organization, accompanied by reduction in repressive marks ^14,18^. Furthermore, the observed central chromatin in the live lamin C OE further supports the ability of our imaging setup to detect chromatin in the interior of the nucleus, when it is indeed present.

### A computational chromatin polymer model to describe the shift from peripheral to central chromatin organization

Computational modeling is a powerful tool to study spatial genome organization and has the potential to suggest mechanistic insights and governing principles of the 3D chromatin organization, based on physical and biological processes and experimental data ^41^. To gain mechanistic insights into the observed shift in nuclear-scale chromatin distribution from the periphery to the interior, we performed simulations of a polymer-based chromatin model.

The computational model approximates chromatin as a semi-flexible, bead-spring polymer ^4,42–44^, and shown schematically in Figure 5A. Each spherical bead has a diameter of 10 nm that accounts for 600 bp of DNA and coarse-grains over 3 nucleosomes ^42^. Neighboring beads along the chain are connected via springs (not shown) and each bead interacts with any other bead in the chain with a repulsive, Lennard-Jones (LJ) potential ^4,42^ that is repulsive at very short range of about one bead diameter and can be attractive in the range of a few bead diameters. We simulated two cases: a case of self-attraction, where the interaction is attractive in the range of one to about two bead diameters, and a case of no self-attraction, where the potential is zero for particle separations of more than one bead diameter ^45^. The former case represents generically “self-attracting” chromatin, without the need to specify the small molecular actors, whereas the latter case results in an “excluded volume random walk” of chromatin in an aqueous solvent. This bead-spring model behaves as a flexible polymer on large length scales. To account for the local rigidity in the polymer, we added angular interactions between beads (bending energies) so that our model polymer has a persistence length of 1.2 kb, consistent with a previous estimate of the persistence length for the interphase chromosome as 1–2 kb ^46^. The overall polymer length included 37,333 beads, representing the size (22.4 Mb) of *Drosophila* chromosome X (ChrX). The nucleus was modeled as a hollow spherical shell with a thickness of 10 nm, and beads that comprise this shell account for the nuclear lamina layer, which interacts attractively with LADs of the chromatin ^15^. The volume fraction of chromatin (and its strongly-bound aqueous/small protein molecules) in the nucleus was set to 0.3 ^47,48^.

**Figure 5:**
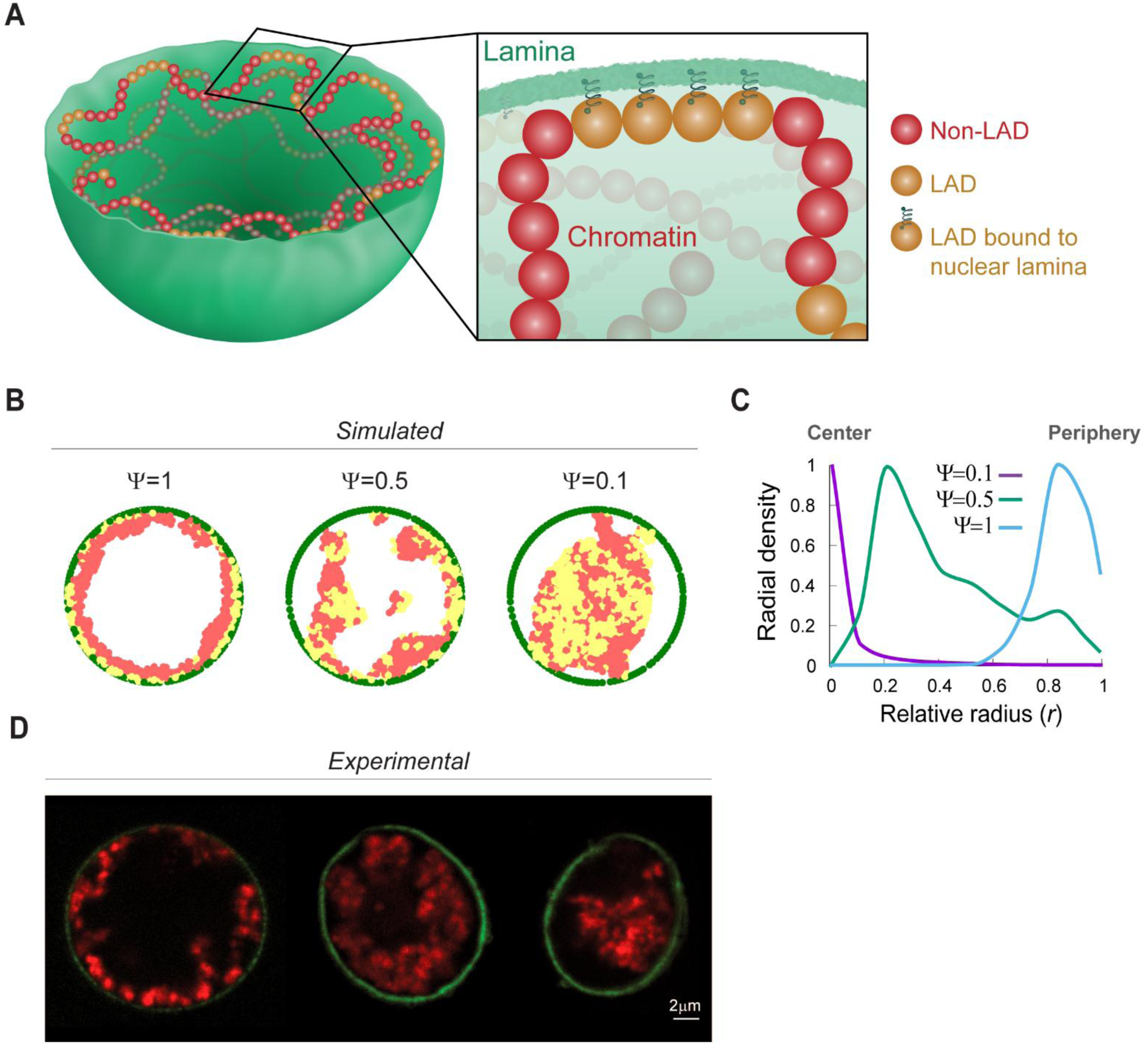
Simulations of 3D chromatin polymer model suggest governing principles for global chromatin organization and its dependence on LAD-nuclear lamina interactions. (A) Chromatin is described by a semi-flexible, bead-spring polymer model that is confined to a sphere, with non-LAD (red) and LAD (yellow) chromatin beads. Yellow LAD beads bound to the lamina are represented with a spring. (B) Simulated results of chromatin concentration maps (LAD in yellow and non-LAD in red) confined within the nuclear lamina (green) for decreasing (left to right) fraction of LADs bound to nuclear lamina (***ψ***). Equatorial plane of the spherical model nucleus is shown. (C) Mean simulated radial chromatin density profiles describe a shift in chromatin distribution, from periphery to the center, with ***ψ*** decrease from 1 to 0.5 to 0.1. (D) Mid XY-planes of experimental live larva muscle nuclei show similar trend in chromatin shift from peripheral distribution (left, control), to more central chromatin distribution in lamin C OE (center and left, intermediate and strong phenotype, respectively).

The polymer comprises two types of beads: lamin-associating and non-lamin-associating (red and yellow in Figure 5A, respectively). When a lamin-associating bead is within a distance of 25 nm of a lamin bead, a strong bond is formed (represented by a spring in Figure 5A). The LADs locations were based on experimentally sequenced data on *Drosophila* ChrX, which also suggests that about 48% of the chromatin consists of LADs ^49^. While we have used numerical values for relevant variables such as bead size, chain length etc., the general trends shown below did not depend on these details and were mostly sensitive to the values of chromatin volume fraction and the fraction of LAD domains interacting with the lamina, as discussed below.

Simulations were performed with coarse-grained molecular dynamics using LAMMPS software ^50^. When the simulation begins, all the chromatin beads are distributed randomly in the interior of the lamin confined sphere (Movie S5). To model changes in the interactions of LADs with the nuclear lamina due to genetic changes of lamin proteins, as well as different fractions of LADs that may be applicable to chromosomes in other organisms, we defined a parameter ***ψ*** as the fraction of LAD beads in chromatin that can form bonds with the nuclear lamina beads. Experimental observations upon lamin C OE suggest weaker chromatin-nuclear lamina interactions. We thus allowed for the possibility that in some systems, not all LADs form bonds with the nuclear lamina, by considering general values of **0 < *ψ*** ≤ **1**.

Figure 5B shows typical simulated results of the chromatin concentration map (yellow for LAD and red for non-LAD beads), confined within the nuclear lamina (green), in the equatorial plane of the spherical model nucleus. The fraction of bound LADs, ***ψ***, decreases from left to right, from ***ψ***=1 representing all estimated 48% LADs in ChrX as bound, to ***ψ***=0.5 and ***ψ***=0.1. Decreased fraction of bound LADs resulted in a shift in the peak average radial chromatin density profile, from the periphery towards the center (Figure 5C). In parallel, Figure 5D shows the experimental mid-XY planes of nuclei from control larva muscle (left), lamin C OE with intermediate phenotype (middle), and lamin C OE with strong phenotype (right). Interestingly, the simulated trends in chromatin concentration profiles and their central shift with decreased LAD binding, closely resemble the experimental results.

While the simulated results are not designed to mimic the details of the experiments, the ability to achieve peripheral chromatin organization at equilibrium, with reasonable biophysical parameters, suggests possible governing principles for 3D nuclear-scale chromatin organization in live nuclei related to the competition between lamin attraction, confinement and polymer entropy and self-attraction of the chromatin. Moreover, the qualitative agreement of the simulations shown in Fig. 5 with the experiments suggests that self-attraction must be included in the model. If the chromatin interactions were purely repulsive (excluding the volume of beads that are close in 3D space), the chromatin would uniformly fill the nucleus for volume fractions of 0.3 (Figure S3). The peripheral chromatin distribution in live control nuclei might therefore arise from strong chromatin interaction with the nuclear lamina, coupled with high chromatin-chromatin affinity (Figure 5A). When LAD-nuclear lamina interactions are decreased, the balance is shifted and the strong chromatin-chromatin interaction gradually drives chromatin towards the center of the nucleus.

## Discussion

Robust chromatin organization within the nucleus has been linked to the control of gene transcription ^41^, yet, information on the three-dimensional chromatin distribution under physiological conditions in a living organism is limited. Here we have revealed the 3D distribution of chromatin in muscle nuclei of live *Drosophila* larvae at high resolution. Our analysis demonstrates a novel mode of chromatin organization at the nuclear mesoscale of fully differentiated cells, wherein a high density of chromatin, including both active and repressed regions, is found at the nuclear periphery. Chromatin density decreases sharply as a function of distance to the center of nucleus, giving rise to a chromatin-devoid region at the center of the nucleus. Importantly, we show that visualization of this new mode of nuclear-scale chromatin architecture was possible primarily due to our ability to image muscle nuclei in unfixed tissue within a live organism, where nuclear volume was preserved. We show that the peripheral chromatin organization was sensitive to lamin A/C levels, and further provide physical insight into these observations by modeling chromatin-LADs interactions. Model simulations show that a peripheral chromatin organization was obtained when LADs and chromatin attractive interactions dominate the entropic tendency to distribute the chromatin uniformly throughout the nucleus. Taken together, our results suggest a novel mode of nuclear architecture with organization into peripheral-chromatin-containing, and central-chromatin-devoid regions in non-dividing, fully differentiated cells, which may impact gene regulation.

A key experimental feature that allowed us to view nuclear partitioning was preservation of nuclear volume in the intact live larval muscles. Previous global chromatin organization studies were performed primarily on fixed preparation ^6,35^, or on cultured cells ^2,30,31^, each causing experimental artifacts that might obscure the native spatial chromatin distribution. Prevailing fixation methods may cause cell and nuclear volume changes ^8,25^, giving rise to shape changes mainly in the Z-direction ^51^, while the dimensions in the imaging plane are less affected, due to cell adherence to the surface and nuclear-cytoskeletal interactions. Furthermore, common fixation reagents can induce significant structural changes to the various cellular and nuclear components, including DNA and RNA ^36,52^. In live cell culture, nuclear and cytoplasmic volumes were shown to be interconnected and to adapt to external signals, through tight control of their mutual water content ^24,26,53^. Cells grown on rigid surfaces, and their nuclei, are relatively spread and flat, with 40%-50% less water content relative to cells grown on soft matrices ^25,26^. Significantly, changes in nuclear volume and shape were shown to have critical functional effects such as altered chromatin organization ^23^, gene expression ^54,55^, and DNA synthesis ^53^. Although in some carefully designed cell culture studies it was shown that nuclear morphology and global chromatin organization are largely preserved upon fixation ^2,36^, both conditions are far from their native tissue environment. Thus, the sensitivity and adaptability of the nucleus and the chromatin to their environment requires careful interpretation of the results of nuclear architecture studies. In view of that, the ability to evaluate nuclear organization in physiologically relevant conditions, as offered by our experimental setup, is of critical importance.

Recent study challenged the prevailing view that repressed and active chromatin compartments partition radially within the nuclear volume, i.e., repressed chromatin predominates at the nuclear periphery and active chromatin prevails near the nuclear center ^35^. In agreement with this report, we did not observe a clear radial separation between active and repressed chromatin distributions, and active chromatin was also detected at the nuclear peripheral compartment.

Interestingly, we observed nuclear partitioning into a peripheral high-density chromatin layer and a central region that is devoid of chromatin not only in muscle fibers, but also in larval epidermal nuclei, implying that this nuclear organization is not unique to muscle fibers. Notably, the muscle and the epidermal cells contain relatively large nuclei, which are also polyploid, thus facilitating depiction of the chromatin and the chromatin-devoid regions. Whereas our findings may represent a specific phenomenon relevant to fully differentiated *Drosophila* larva polyploid tissues, the observation of similar peripheral chromatin organization in 8-cell stage IVF bovine embryo further supports a broader relevance of our findings ^10^. Moreover, in agreement with the sensitivity of chromatin organization to the nuclear volume observed in our study, Popken et al. reported a shift from peripheral to dispersed chromatin architecture that correlated with a reduction in the nuclear volume and with embryonic stage in fixed tissue ^10^. Future analysis of live proliferating smaller nuclei, e.g. in the embryo, or in tissues of non-*Drosophila* organisms, will be required to confirm the generalization of our observations.

The nuclear lamina controls radial chromatin organization by its dynamic tethering of chromatin to the nuclear periphery through LADs ^13,14^. Since the nucleus is thought to be a mechanosensitive organelle ^56–58^, in which the close proximity of the chromatin to the nuclear lamina might sensitize the chromatin to both mechanical inputs, as well as to changes in the nuclear lamina composition, a peripheral chromatin organization might couple between mechanical inputs and transcriptional regulation. This is especially relevant for fully differentiated non-dividing cells, in which the program of gene transcription has been well established. In addition, the global chromatin detachment from the periphery and its condensation towards the center that was observed upon elevated lamin C levels agrees with previous reports on reduced chromatin-lamina tethering following up-or down-regulation of lamin C, as well as with observations in cells with laminopathy-associated mutations. Interestingly, such chromatin structural changes were accompanied by chromatin de-condensation and reduction in the repressive HP1 and H3K9me3 marks ^15–19^. Further investigation into specific epigenetic and transcriptional alterations is required to appreciate the functional significance of such substantial changes in global chromatin organization in the muscle nuclei.

The dominant effect of lamin C over-expression that we detected might result from masking of chromatin binding sites to LEM-domain proteins, thus lowering the probability of chromatin association with the lamina through LADs and driving overall condensation towards the nucleus center due to chromatin self-attraction. Alternatively, chromatin condensation away from the periphery might be driven by increased chromatin cross-linking with lamin C in the nucleoplasm, as suggested previously ^59^.

The simulations of chromatin distribution dynamics involved only a small number of parameters extracted from the literature, including the percentage of LADs per chromosome, the tendency of chromatin to effectively self-attract ^45^, and the extent of nuclear confinement. The similarity between the simulation and the experimental results suggests that these are among the most critical factors in determination of the partitioning of chromatin on the nuclear scale. It further suggests that a balance between chromatin-lamina interactions through LADs, entropy, and chromatin self-association may represent the major driving forces for the nuclear partitioning observed experimentally. The model has also provided us with insights for future experimental tests, for example by changing the nuclear lamina properties.

In summary, our study reveals a novel mode of nuclear-scale chromatin organization in fully differentiated cells, in which chromatin density is high at the nuclear periphery and low in the nuclear center, with an effectively central chromatin-devoid region. This could only be observed in live nuclei, within their native physiological tissue environment, since fixation can lead to dehydration, which either alters chromatin organization or obscures its actual 3D visualization. Simulation of chromatin organization based on LADs binding and chromatin self-attraction recapitulated the experimental observations, and predicted that changing each of these parameters may disrupt peripheral chromatin organization and change nuclear-scale chromatin density profile. Indeed, over-expression of lamin C in our experiments disrupted the peripheral chromatin organization, supporting the basic assumptions of the model. Future experiments should address extrapolation of these findings to other cell types and animal models, as well as the functional consequences of alterations in global chromatin organization.

## Materials and Methods

### Fly stocks and handling

The following fly stocks were used: *ubi-H2B-mRFP/CyO; Dr/TM6B* ^60^, *CyO/Sp; UAS-laminC-GFP* ^40^, *UASp-mintbody-EGFP; Pr,Dr/TM3,Sb,Ser* ^38^, *His2Av-GFP* (FBst0005941), *GAL4-Mef2*.*R* (FBst0027390), UAS-mCherry-NLS (FBst0038424), UAS-klar-GFP ^61^. All crosses were carried out at 25°C and raised on cornmeal agar.

### The minimal constraint device for imaging live intact Drosophila larvae

For imaging live nuclei in their intrinsic environment, a minimal constraint device for *Drosophila* larvae was designed in our lab, to be placed on top of a confocal microscope stage, as previously described ^29^. Shortly, the center of the device is a thin plastic bar positioned in a larger frame, with a groove crossing the frame and bar from side to side. The larva is placed in the groove, and glass capillaries (Drummond Scientific Company, PA, USA, cat. #1-000-0010, O.D 0.026”, I. D. 0.0079”), which are aligned with the larval body, are glued to its head and tail (Superglue liquid, Pattex, Australia). The device base with a coverslip glass in it are placed on top of the larva in the bar, and all the parts of the system are inverted together as a unit. To keep the larva moist during the experiment, it is enclosed with alginate hydrogel made by polymerizing a solution of 4% alginate (cat. # 180947, Aldrich) dissolved in 0.9% saline (NaCl, cat. # 0277, J. T. Baker) with a solution of 0.8 M of CaCl_2_·2H_2_O (cat. # 2382.0.0500, Merck).

### Live preparations

Third instar, wandering larvae were selected for imaging. Prior to placement of the larvae in the device, it was immersed in water for ∼3 hours to decrease its movement (larval movement could be restored by exposure to air). For each larva, at least 3 nuclei were imaged from randomly chosen muscles along the entire larval body. Before and after obtaining full Z-stacks of the nucleus, a single middle XY and middle YZ planes were taken to verify that the nucleus did not deform or move during image acquisition.

### Fixed preparations

Third instar, wandering larvae were selected and dissected into flat preparations as previously described ^61^. Paraformaldehyde (4% from 16% stock of EM grade, 15710, Electron Microscopy Sciences) was used for fixation. Specimens were fixed for 20 minutes, washed several times in PBS, and mounted in Shandon ImmunoMount (Thermo Fischer Scientific).

### Microscopy and image acquisition

Confocal imaging of the live and fixed preparations was performed using an inverted Leica SP8 STED3X microscope, equipped with internal Hybrid (HyD) detectors and Acusto Optical Tunable Filter (Leica microsystems CMS GmbH, Germany) and a white light laser (WLL) excitation laser. Spectral analysis of His2B-RFP and klar-GFP channels revealed their maximal emission to be at excitation wavelengths of 586 nm and 478 nm, respectively. RFP emission signal was collected at the range of 597 – 699 nm and GFP emission in a range of 488 – 559 nm. All nuclei were imaged with a HC PL APO 86x/1.20 water STED White objective, NA=1.2, at a scan speed of 400 Hz and a pinhole of 0.8 AU. Z-stacks were acquired using the galvo stage, with 0.308 µm intervals. Bit depth was 12 and to enhance image quality, field of view (FOV) and laser intensity were adjusted separately for each nucleus sampled. The acquired images were visualized during experiments using LASX software (Leica Application Suite XLeica microsystems CMS GmbH).

### Image processing and data analysis

Only nuclei that did not move or deform during image acquisition were included. Arivis Vision4D 3.1.2-3.1.4 was used for image visualization and analysis. Nuclei labelled for nuclear membrane (control and lamin C-GFP over-expression groups) were segmented in 3D with a dedicated pipeline based on the ‘blob finder’ operation. For nuclei co-labeled with His2B-mRFP and H3K9ac-mintbody-EGFP, a separate pipeline was used for segmentation, based on automatic ‘otsu’ threshold operation, with segment modification to close a 3D nucleus object. All segmented nuclei were further divided into 10 concentric 3D shells (Figure 1A), using a dedicated Python script embedded within the Arivis Vision4D environment. Cytoplasmic background intensity was calculated from additional shell generated outside of the segmented nucleus using the ‘dilate’ segment modification operation. Nuclei and corresponding shell segment volumes, dimensions, and channel intensities were then exported to MATLAB R2019b (MathWorks) to generate radial distribution profiles. Chromatin density for each shell was calculated by summing the His2B fluorescence intensity within the shell and dividing it by shell volume. To compare radial profiles of nuclei from different muscles and larvae, the absolute His2B density values were divided by His2B density in the cytoplasmic shell.

### Statistics

For each group we sampled at least 3 nuclei per larva, and 5 different larvae were analyzed. To compare radial distribution between groups, each radial distribution profile of an individual nucleus was divided by its maximum value. The nuclei were fitted with a 3^rd^ order polynomial, using a linear mixed effects model fit by REML, accounting for group as a fixed effect, and for nucleus and larva as random effects. Analyses were done using the packages ‘lme4’ and ‘lmerTest’ in R v.3.5.1.

## Acknowledgments

We thank Kimura Hiroshi (Osaka University, Japan), Kazuhiro Furukawa (Niigata University, Japan), and the Bloomington Stock Center for providing fly lines. The images in this paper were acquired at the Advanced Optical Imaging Unit, de Picciotto-Lesser Cell Observatory unit, at the Moross Integrated Cancer Center Life Science Core Facilities, Weizmann Institute of Science. We thank Michal Shemesh and Yosef Addadi for advising on image acquisition, Ofra Golani for advising on imaging analysis and Ron Rotkopf for statistical analysis, all from the Life Sciences Core Facility, Weizmann Institute. Special thanks to Maurizio Abbate from Arivis V4D support team, for incorporating custom Python script for 3D radial shells. We are grateful to Georges Ankaoua, and Benjamin Pasmantirer from the Physics Core Facilities at Weizmann Institute for helping in the design of the live imaging device. This study was supported by grants from “The French Muscular Dystrophy Association (AFM-Téléthon)” grant # 22339, NSF-BSF (BSF grant # 2016738), Israel Science Foundation (ISF) grant # 750/17, and the Weizmann - Krenter-Katz - Interdisciplinary Research at the Interfaces of Life and Exact Sciences.

