## Supplementary Information for "Live imaging of chromatin distribution in muscle nuclei reveals novel principles of nuclear architecture and chromatin compartmentalization" (from another version of this preprint: version 1, 2020-06-21)

### Supplementary Movie S1

Live, *in-vivo,* Drosophila larva muscle nucleus exhibiting partitioning into peripheral chromatin layer and central chromatin-devoid compartment.
Nuclear envelope is labeled with klar/nesprin-GFP and chromatin is labeled with His2B-mRFP. The nucleus is positioned on top of the sarcomeres (white), observed by fluorescence reflection mode. 3D reconstructed image from confocal z-stack obtained at every 0.3 µm.

### Supplementary Figure S1

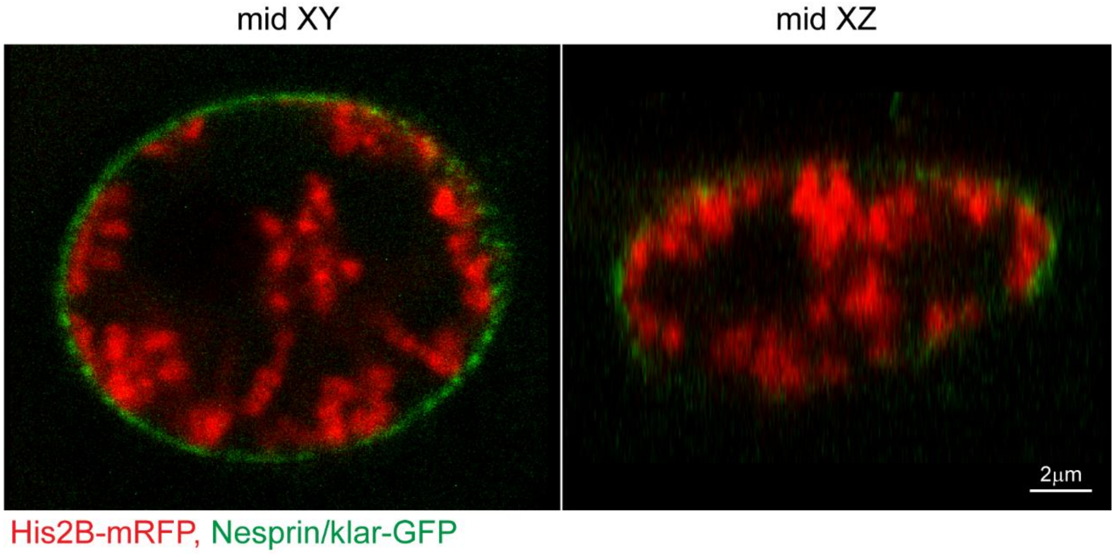


Live muscle nucleus with peripheral chromatin organization but high His2B-mRFP density at the center caused by chromatin filament extended between two sides of the nucleus.
Mid XY (left) and XZ (right) planes of confocal z-stack.

**Supplementary Figure S2**

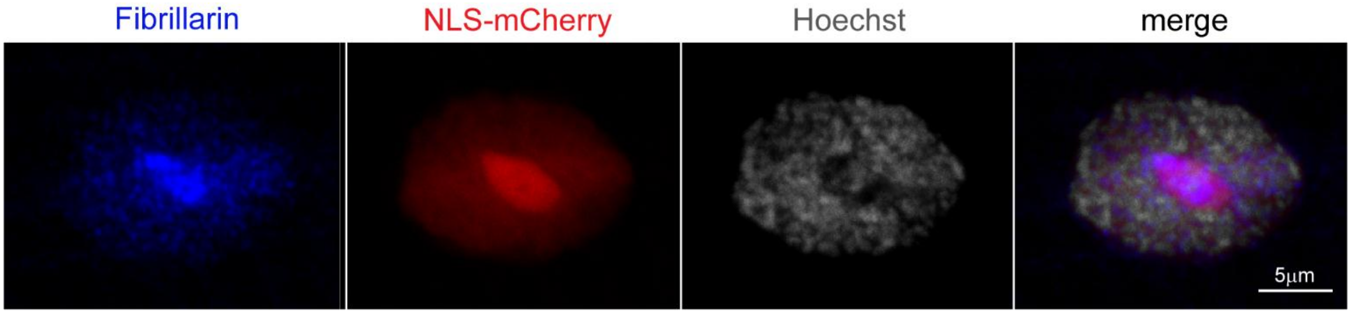


NLS-mCherry shows denser accumulation at the nucleolus.
Maximum intensity projection of representative muscle nucleus from fixed, flat larva preparation. Nucleolus is labeled with anti-fibrillarin (blue), and overlaps with dense NLS-mCherry labeling within the nucleus (red). DNA is labeled with Hoechst (white).

**Supplementary Movie S2**

Live epidermal nucleus from *Drosophila* larva, labeled with His2B-mRFP, demonstrates a peripheral chromatin distribution. 3D reconstructed image from confocal z-stack.

**Supplementary Movie S3**

Active chromatin regions are distributed within the peripheral chromatin layer in live muscle nucleus.
Active chromatin regions labeled with H3K9ac-EGFP (green), and total chromatin labeled with His2B-mRFP (red). Active, diffused chromatin is not identified within the chromatin-devoid compartment at the center of the nucleus. 3D reconstructed image from confocal z-stack obtained at every 0.3 µm.

**Supplementary Movie S4**

Lamin A/C over-expression in the muscle causes chromatin detachment from the nuclear lamina, and condensation towards the center.
Nuclear envelope is labeled with UAS-LamC-GFP (green), and total chromatin with His2B-mRFP (red). 3D reconstructed image from confocal z-stack obtained at every 0.3 µm.

**Supplementary Movie S5**

Simulation movie of polymer-based model of chromatin. Mid plane of the spherical model nucleus. Lamin beads in green, chromatin LAD beads in yellow, and chromatin non-LAD beads in red. Simulation start with all beads randomly distributed in the center of the nucleus. First 10 seconds represent chromatin with no self-attraction property, leading to yellow LAD binding to the lamina and uniform distribution of the remaining chromatin beads, within the entire nuclear volume. At t=10 sec, chromatin self-attraction is turned on, resulting in rapid reorganization of all chromatin beads into a peripheral chromatin layer.

**Supplementary Figure S3**

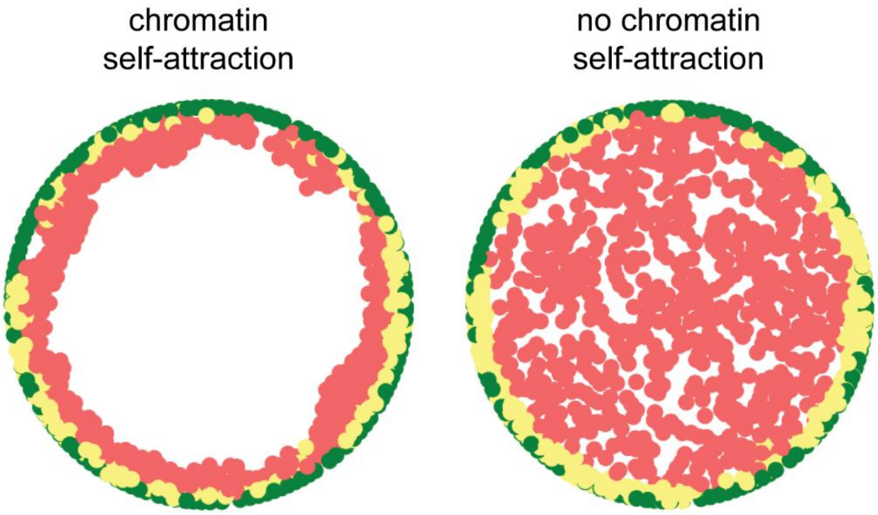


Peripheral chromatin organization is driven by chromatin attachment to the lamina through LADs, coupled with chromatin self-attractive property.
Representative mid plane of the spherical model nucleus. Lamin beads in green, chromatin LAD beads in yellow, and chromatin non-LAD beads in red. Incorporating chromatin self-attraction during simulation results in peripheral chromatin organization at steady state (left) versus uniform chromatin distribution, when chromatin is not self-attractive (right).
